# Movement of moths through riparian reserves within oil palm plantations

**DOI:** 10.1101/204990

**Authors:** Ross E. J. Gray, Eleanor M. Slade, Arthur Y. C. Chung, Owen T. Lewis

## Abstract

Tropical forests are becoming increasingly fragmented through conversion to agriculture, with negative consequences for biodiversity. Movement of individuals among the remaining fragments is critical to allow populations of forest-dependent taxa to persist. In SE Asia, conversion of forests to oil palm plantations is a particular threat. Many oil palm dominated landscapes retain forested riparian reserves along streams and rivers, but the extent to which these riparian reserves are used for movement, relative to the oil palm matrix, is poorly understood. We used mark-release-recapture methods to investigate the movement of five moth taxa in riparian reserves connected to a continuous forest area within oil palm matrix in Sabah, Malaysian Borneo. Moths were recaptured on average 68m from the release point, and the mean time to recapture was between 2 and 3 days. Moths showed a significant tendency to move within forested riparian reserves than into adjacent oil palm. When moving within riparian reserves moths also showed a tendency to orient their movement away from the continuous forest into the riparian reserve rather than towards continuous forest or into oil palm. Overall, our results show a role for riparian reserves as movement corridors through oil palm plantations for some invertebrate species, strengthening the case for their retention and re-establishment.

## INTRODUCTION

Tropical forests are increasingly degraded, fragmented and converted to agriculture, with detrimental effects on biodiversity (Gardner et al., 2009; Barlow et al., 2010; Hansen et al., 2013; Haddad et al., 2015). South East Asian forests have been particularly affected, with the Indo-Malayan biodiversity hotspot experiencing pronounced biodiversity loss (Myers et al., 2000; Sodhi et al., 2004; Sodhi et al., 2010; Gibson et al., 2011; Ewers et al., 2015; Lewis et al., 2015). A major threat is the rapid expansion of areas planted with oil palm (*Elaeis guineensis* Jacq.) (Fitzherbert et al., 2008; Gaveau et al., 2014), which is now the world’s primary source of vegetable oil and fat (Turner et al., 2008). Oil palm is thus a vital part of the economies of Southeast Asia, accounting for 3.82% of the gross domestic product in Malaysia alone (∼$896 million; Mahidin, 2018). As mosaic landscapes that incorporate both natural forest and oil palm agriculture become increasingly common, it is important to understand how best to manage and design these landscapes to support their remaining biodiversity.

Frequent features of mosaic oil palm landscapes are riparian reserves (also called buffer zones or riparian strips)(Luke et al., 2019). These are linear areas of riverine forest, set aside primarily to reduce run-off into streams (Sweeney et al., 2004). Riparian reserves improve water quality (Mayer et al., 2007) and benefit aquatic and forest-dependent terrestrial fauna (Ricketts, 2004; Marczak et al., 2010; Gray et al., 2014). They may also serve as movement corridors linking fragments to continuous forest (Beier and Noss, 1998; Tewksbury et al., 2002). Recently, riparian reserves have been added as a requirement for Roundtable on Sustainable Palm Oil (RSPO) certification, with a minimum forest buffer of 30 to 100 m on both sides of the river, with the minimum width stipulated depending on river width, reserve placement, and perceived use (Barclay et al., 2017; Lucey et al., 2017).

As rainforest habitats become increasingly fragmented and isolated, connectivity between forest fragments may become critical to allow populations to persist (Hanski, 1999; Ewers and Didham, 2006; Lucey and Hill, 2012). Understanding the movement ability of flora and fauna is therefore a key consideration in conservation and management strategies in human-modified landscapes. Movement ability is influenced by the behavioural responses of different species to habitat boundaries (Lucey and Hill, 2012; Kallioniemi et al., 2014), the physical costs of movement (Bonte et al., 2012), and the permeability of the matrix (Ewers and Didham, 2006; Scriven et al., 2017). Forest dependent-taxa (i.e., those that need forest to support viable populations) are the most vulnerable to fragmentation, as their restricted ranges and reluctance to cross forest to non-forest boundaries result in small, isolated populations which suffer local extinctions with little prospect of recolonization (Sodhi et al., 2010; Scriven et al., 2015). However, only a few studies have investigated the movement behaviour of tropical forest-associated taxa (Bouchard and Brooks, 2004; Brouwers and Newton, 2009; Lucey and Hill, 2012; Khazan, 2014; Scriven et al., 2017).

Here, we examine the movement behaviour of forest-associated moths (Lepidoptera) in riparian reserves within oil palm landscapes in Sabah, Malaysian Borneo. Moths are a species-rich and functionally important insect group in tropical forests, acting as pollinators, herbivores, and prey for a wide variety of vertebrate and invertebrate taxa (Kitching et al., 2000; Summerville et al., 2004; Slade et al., 2013). Tropical forest moths are relatively understudied in terms of dispersal and movement. Only one study to our knowledge explicitly measures movement capacity in tropical moths, assessing the distance to which moths are attracted to a light trap (Beck and Linsenmair, 2006), with the closest comparable study grouping moths by dispersal classes and measuring the correlation of these classes with certain dispersal traits (Beck and Kitching, 2007). Studies in temperate agricultural landscapes have shown that habitat connectivity is critical for moth movement (Merckx et al., 2010; Slade et al., 2013).

To understand the potential role of riparian reserves as movement corridors for tropical moths, we used mark-release-recapture methods, a common approach for the study of animal movement (e.g. Lewis et al., 1997; Hanski, 1999; Slade et al., 2013). We investigated (1) whether forest-associated moths were more likely to move through riparian reserves than into oil palm; and (2) whether there was tendency for these moths within riparian reserves to move towards or away from the adjacent areas of continuous forest to which the riparian reserve is connected. In combination, these analyses allow us to compare the likely permeability of oil palm plantations and riparian reserves for moths, and their potential utility in linking forest fragments.

## MATERIALS AND METHODS

### STUDY SITES

Fieldwork took place between November 2016 and April 2017 within the Stability of Altered Forest Ecosystems (SAFE) project landscape in South Eastern Sabah, Malaysia (4°38” N to 4°41” N, 117°31” E; Ewers *et al*. 2011; Fig. 1A). We selected three focal sites where a riparian forest reserve, embedded within an oil palm matrix, was connected to a large block of continuous virgin jungle reserve (VJR) of lowland dipterocarp rainforest. The VJR covers 2,200 ha of land and is connected to a larger (> 1 million ha) area of protected forest (Ewers et al., 2011). The oil palm plantations at each site were approximately the same age (∼8 years). The three focal sites (RR3, RR10, RR18) were selected to be similar in configuration and structure and, in each case, had tall trees (some >40m), high canopy cover, and similar mean riparian forest widths (48m, 58m and 41m in RR3, RR10, and RR18 respectively (calculated from satellite imagery of the sites (M. Pfeiffer, unpublished data)). However, there were inevitably minor variations among sites in the precise configuration of landscape elements (Fig. S1).

**Figure 1.**
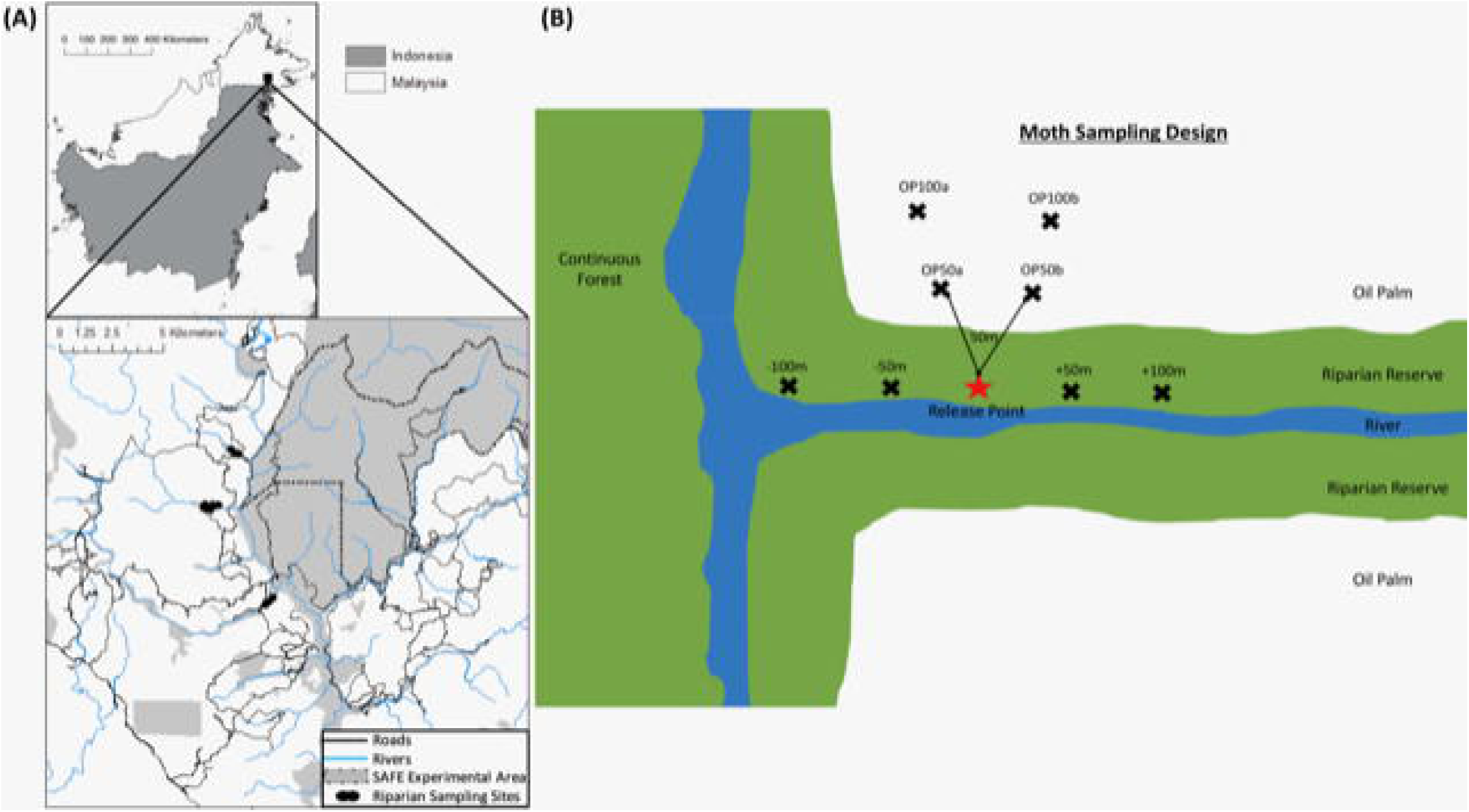
**(A)** The upper panel highlights the location of the study area in Sabah, Northern Borneo. The lower panel displays a map of riparian reserve sites used in this study. The map is adapted from Gray et al. (2014). Black dots represent the riparian sampling sites labelled RR18, RR3 and RR10 from top to bottom. The dotted black line shows the SAFE project experimental boundary. The solid grey shaded area in the top panel is the remaining logged forest, blue lines indicate rivers, and the white area indicates the matrix of oil palm plantations. **(B)** Schematic representation of the general sampling design in each riparian reserve for the mark-release-recapture study for moths. Black crosses indicate trapping points which were spaced by at least 50 m. Lines on the diagram are to indicate spacing and are not drawn to scale. The star indicates the release point of marked moths.

### SPECIES SELECTION

Pilot studies identified five forest-associated moth taxa which could be trapped in sufficient numbers for a focused study: *Erebus ephesperis* Hübner, *Erebus camprimulgus* Fabricius, *Ischyja* spp. Hübner, *Erebus gemmans* Guenée and *Hypoprya* spp. Guenée (Holloway, 2005). Multiple *Ischyja* and *Hypoprya* species were present in our samples; living individuals could not always be identified reliably to species in the field, so individuals within each of these genera were pooled for analyses. The focal moth taxa are nocturnal, relatively large-bodied and common within the focal landscape. Their larval hostplants – essential to maintain populations – are woody vines, shrubs and trees (Holloway, 2005) which are absent from oil palm plantations.

### MOTH MOVEMENT

Moth movement was studied by releasing individuals at a single, central point where the riparian reserve connected to the VJR at each site. In each of the three study sites we established a network of baited traps, spaced a minimum of 50m apart, with a single central release point (without a trap) (Fig. 1B; individual site diagrams in Fig. S1). In RR10 and RR18, six traps were placed within oil palm and six traps in the riparian reserve. However, the shape of the riparian reserve allowed only 4 traps to be placed in the oil palm at RR3. We used Bugdorm© Pop-up Butterfly Bait Traps with 10 cm cone openings, baited with fermented banana and suspended so that the base was at least 1 m above the ground. During each round of sampling, traps were set on day 1 and allowed to accumulate moths for 24 hours. On day 2, individuals of the focal moths were removed from the trap sequentially and marked on the wing with a unique number using a black Sharpie™ permanent marker (Betzholtz, 2002; Truxa and Fiedler, 2012). Moths were transported in a ventilated, sealed container to the release point, where they were released into dense vegetation while inactive. Data on species, trap of capture, release date, and recapture data were recorded for each individual. Baits were replaced each day and any moths from non-focal species were removed from the traps without marking. On day 3, the abundance of focal moths in the traps was recorded, any new moths were marked and released from the release point, and any recaptures were recorded. The protocol for day 3 was repeated until a minimum of 60 recapture events (across all focal taxa) had been accumulated at each site (between 8 to 14 days).

### DATA ANALYSIS

We standardized comparisons by analyzing moth recapture data from a subset of the traps at each site (8 traps per site: 4 in riparian, 4 in oil palm) spaced at comparable distances from the release point (=<100m). We first used a one-tailed binomial test to investigate whether the proportion of moth recaptures in riparian forest was significanty different from that expected under the null hypothesis of random movement (50% expected proportion) to either riparian forest or oil palm. Similarly, for the subset of moths recaptured within the riparian reserves (4 traps per site), we used a one-tailed binomial test to investigate whether there was a tendency for these moths to move towards continuous forest, as opposed to away from it down the riparian reserve. Statistical analyses were carried out using R (R Development Core Team, 2017).

## RESULTS

Across all species and sites, we marked and released 829 moths, of which 281 (33.7%) were recaptured (Table S1). *Hypoprya* spp. had the highest percentage of recaptures (45.5%) and *Ischyja* spp. the lowest (29.6%; Table S1). Of recaptured individuals, 37% were recaptured within 24 hours of release, and the remainder after multiple days (2 to 12 days). Moths moved on average 67.5m over an average of 2.63 days, with distances varying among taxa and among sites (Table S2).

A significantly higher proportion of moth recaptures was recorded in riparian reserve traps than in oil palm plantation traps at comparable distances (N = 283, Expected Proportion = 0.5, Empirical Proportion = 0.622, CI = [0.572, 1.00], *P* < 0.001; Table 1; Fig. 2A). For those moths moving within riparian corridors, individuals were significantly more likely to move along the riparian reserve away from continuous forest than they were to move towards it (N = 176, Expected Proportion = 0.5, Empirical Proportion = 0.665, CI = [0.602, 1.00], *P* < 0.001; Table 1; Fig. 2B).

**Table 1.**
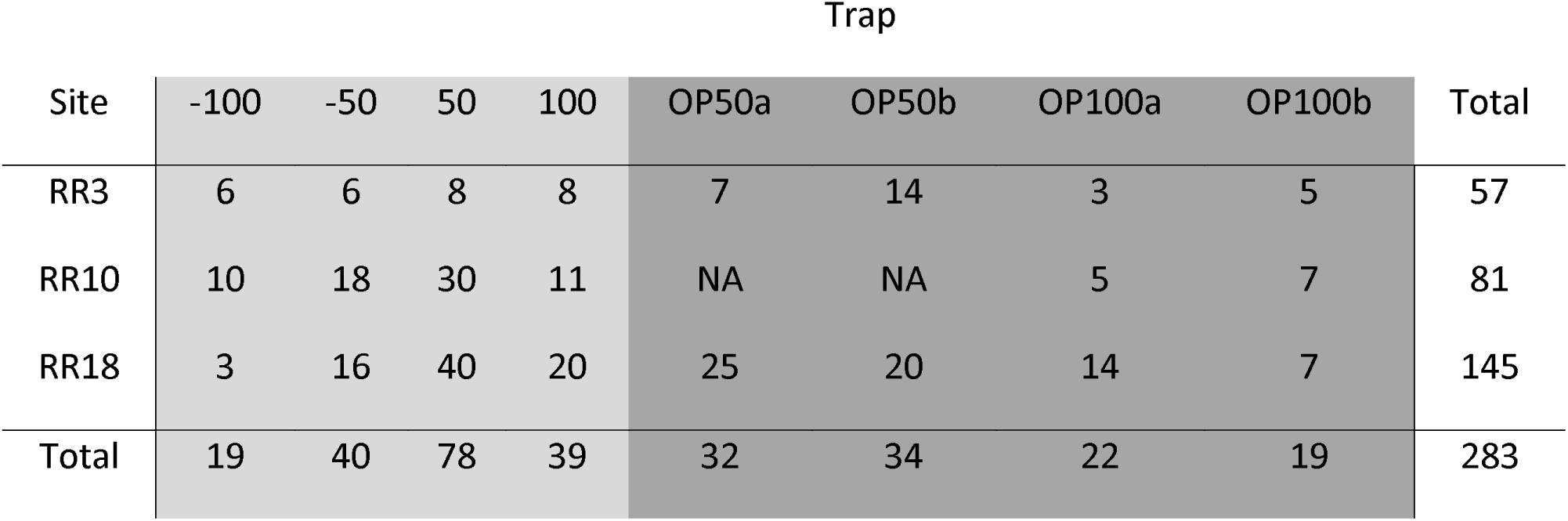
Summary table showing the number of recaptures for each trap in each site. Traps in light grey shading are located in the riparian reserve and traps in dark grey shading are located in the oil palm plantation

**Figure 2.**
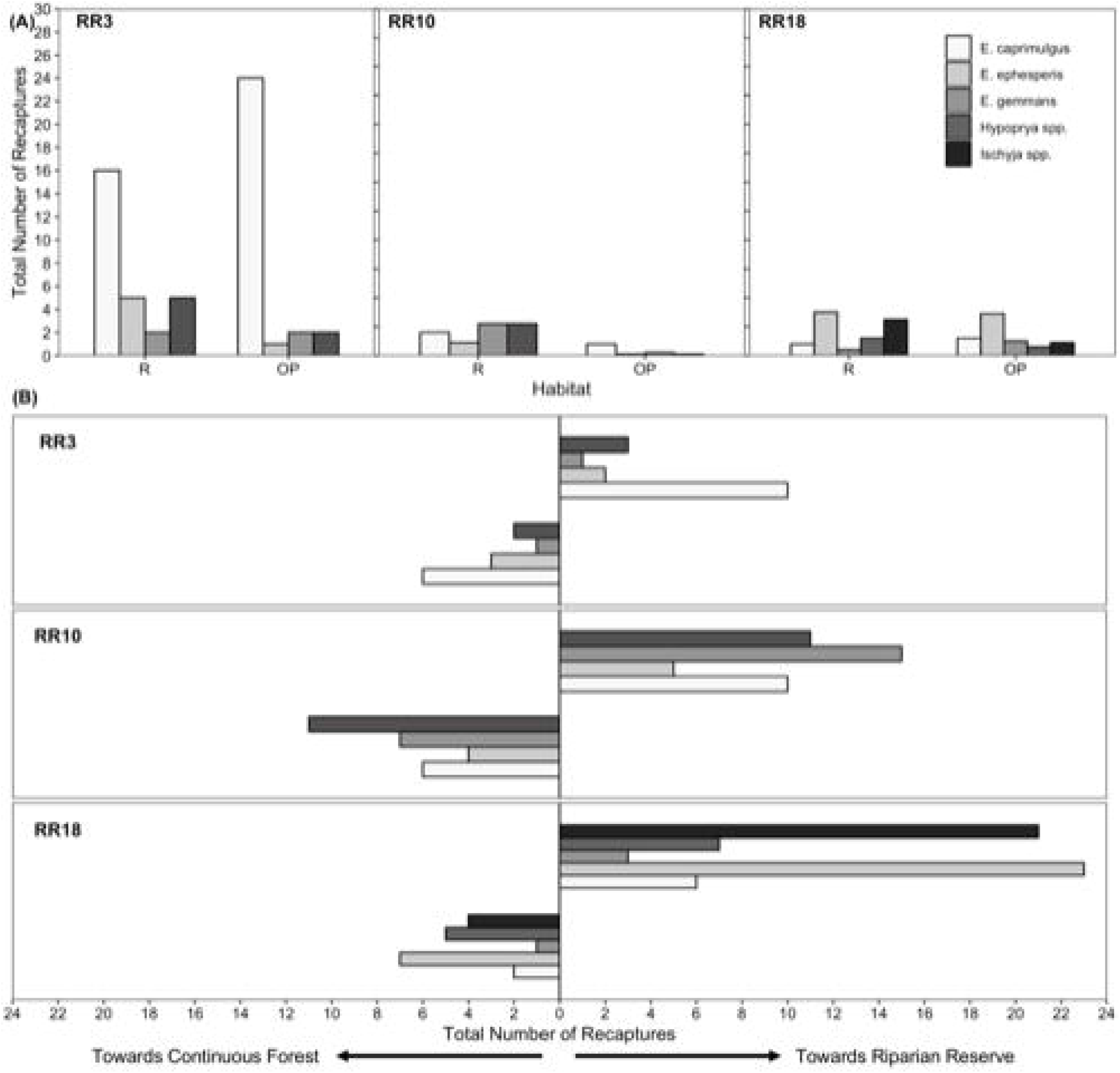
Number of recaptures and orientation of movement for moths. **(A)** Total number of recaptures for each species in each habitat. **(B)** Total number of movement orientations for each species towards riparian forest or towards continuous forest (VJR) in each site. Bars are shaded based on taxon.

## DISCUSSION

Riparian forest reserves can represent important habitat for many invertebrate taxa (Barlow et al., 2010; Gray et al., 2016; Luke et al., 2017; Scriven et al., 2017) but their importance for invertebrate movement is poorly studied. Previous studies on the dispersal and movement behaviour of insects have shown that habitat boundaries are often barriers for forest-dependent species (Gray et al., 2016; Scriven et al., 2017), suggesting that agricultural habitats bordering forest habitats may limit movement (Arellano et al., 2008; Merckx et al., 2010; Slade et al., 2013). Our data provide new information on the movement behaviour of moths in riparian forest corridors embedded within an oil palm matrix.

Movement by the focal moth species responded strongly to habitat type and boundaries. Moths were more likely to be recaptured in traps within riparian forest than in traps in oil palm at equivalent distances. This supports previous studies in temperate regions showing that moth movements are often influenced by habitat boundaries (Betzholtz, 2002; Slade et al., 2013). The moth species investigated are relatively specialised in terms of their larval host plants (Holloway, 2005). The absence of these resources within plantations, along with a contrasting vegetation structure characterised by a more open canopy and associated differences in microclimate, may have discouraged boundary-crossing into adjacent oil palm (Barlow et al., 2010).

Since oil palm traps were placed relatively close to the riparian reserve (between 50 – 150 m), movements of some individuals into the oil palm may represent a ‘spillover effect’, as has been documented for butterflies (Lucey and Hill, 2012; Scriven et al., 2017), ants, bush crickets, beetles and some small mammals in forest-oil palm matrices (Nurdiansyah et al., 2016). Traps were baited with fermenting banana which may be detectable by foraging adult moths over distances spanning the scale of the study. Thus, some moth movements into the oil palm may represent foraging activities for moths that are otherwise dependent on forested areas to maintain viable populations (Scriven et al., 2017). It seems likely that plantations provide feeding opportunities for adult moths in the form of abundant oil palm fruit, so such movements may be typical of these landscapes. However, whether these fruits benefit moth populations in the long term by increasing population growth rates or carrying capacities is unknown.

Our data allow us to assess directionality of moth movements with respect to nearby areas of continuous forest. This is of interest because it allows us to understand whether moths that enter riparian reserves from large blocks of forest to which they are connected are likely to continue along these reserves, or to turn back. Surprisingly, there was a significant tendency for moths to move away from connected forests and further along the riparian reserve. It seems unlikely that this tendency reflects an underlying orientation bias or preference (for example, a tendency to move on a particular bearing), since there was no appreciable directional airflow during the study, which can sometimes dictate orientation of movement (Vickers, 2005), and our three riparian sites were oriented at different angles. Orientation could have been influenced by unmeasured factors such as gradients in vegetation structure and composition. Whatever the mechanism, this behaviour seems likely to facilitate successful crossing of the oil palm matrix in situations where continuous riparian corridors link forest fragments within the landscape, as has been suggested for other invertebrate (Lucey and Hill, 2012; Khazan, 2014) and vertebrate taxa (Medina et al., 2007; Gillies and Clair, 2008; Keuroghlian and Eaton, 2008). Further study will be needed to quantify the distances over which such movements can occur, and the extent to which riparian reserves serve to support viable as opposed to sink populations of these and other forest-dependent species.

We did not seek to explore between-species differences or measure life history traits in this study, but traits are likely to explain interspecific differences in movement distances. In temperate moths, life history traits such as wingspan and wing shape, along with the precise habitat affinities of individual species, can significantly influence movement behaviour (Slade et al., 2013).

Despite the limited number of taxa and sites we were able to investigate, our results suggest that movement patterns and behaviours of tropical moths are amenable to study using mark release recapture methods. Additional work is now needed to assess the radius over which moths are attracted to baits and the extent to which this might influence moth movement patterns (Merckx and Slade, 2014). Light traps are the most frequent method used to survey moth communities, and attract higher numbers and diversities of species from a wider range of moth families than fruit-baited traps (Beck and Linsenmair, 2006). However, light traps are rarely suitable for live capture in tropical forests, as the high activity of trapped moths tends to lead to their damage and death (personal observations). We suggest that baited traps like the ones used in our study have considerable potential for studies of moth movement through fragmented landscapes.

## AKNOWLEDGEMENTS

We are grateful to Sabah Biodiversity Council and Sabah Forestry Department for providing permission to carry out the research under access access licence JKM/MBS.1000-2/2 JLD.3(159). We would also like to thank the South East Asian Rainforest Research Partnership (SEARRP), the Stability of Altered Forest Ecosystems (SAFE) project, and research assistants: Lizzie, Loly, Anis, Zul, Sabidi, and Noy. An earlier version of this manuscript was released as a preprint: Gray et al. (2018).

## AUTHOR CONTRIBUTION STATEMENT

OTL and EMS conceived the research idea and secured funding. OTL and REJG designed the study. REJG collected the data, carried out the analyses and led the writing of the manuscript. All authors contributed to editing the manuscript.

## DATA AVALAILABILITY STATEMENT

The data used in this study are archived on the Environmental Information Data Centre (https://doi.org/10.5285/4ca7f1ff-7bc4-4077-bc9c-96be1be3e655) and on Zenodo (doi to be added following acceptance), with access through the SAFE project website (www.safeproject.net, Dataset ID: to be added following acceptance).

