## Supplementary Information for "Movement of moths through riparian reserves within oil palm plantations"

**Supplementary Tables**

Table S1. Summary table showing the number of marked (M) and recaptured (R) moths at each site.

|  | ***Ischyja* spp.** | | | ***Erebus camprimulgus (male)*** | | | | ***Erebus caprimulgus (female)*** | | | ***Erebus gemmans*** | | | ***Hypoprya* spp*.*** | | | ***Erebus ephesperis*** | | | **Totals** | | |
| --- | --- | --- | --- | --- | --- | --- | --- | --- | --- | --- | --- | --- | --- | --- | --- | --- | --- | --- | --- | --- | --- | --- |
| **Site** | **M** | **R** | **R %** | | **M** | **R** | **R %** | **M** | **R** | **R %** | **M** | **R** | **R %** | **M** | **R** | **R %** | **M** | **R** | **R %** | **M** | **R** | **R %** |
| RR3 | 103 | 16 | 15.5 | | 42 | 15 | 35.7 | 128 | 40 | 31.3 | 13 | 4 | 30.8 | NA | NA | NA | 57 | 15 | 26.3 | 343 | 90 | 26.2 |
| RR10 | 76 | 29 | 38.2 | | 38 | 12 | 31.6 | 67 | 28 | 41.8 | 6 | 0 | 0.0 | 31 | 30 | 96.8 | 37 | 13 | 35.1 | 255 | 112 | 43.9 |
| RR18 | 132 | 49 | 37.1 | | 44 | 14 | 31.8 | 35 | 15 | 42.9 | 45 | 18 | 40.0 | 36 | 30 | 83.3 | 184 | 76 | 41.3 | 476 | 202 | 42.4 |
| Total and Mean % | 311 | 94 | 30.2 | | 124 | 41 | 33.1 | 230 | 83 | 36.1 | 64 | 22 | 34.4 | 67 | 60 | 89.6 | 278 | 104 | 37.4 | 1074 | 404 | 37.5 |

Table S2. Summary table showing the mean, median and maximum dispersal distance over a 24-hour period for each moth species at each site. Error is standard deviation.

|  | ***Ischyja* spp.** | | | ***Erebus camprimulgus (male)*** | | |
| --- | --- | --- | --- | --- | --- | --- |
| **Site** | **Mean Movement Distance** | **Median Movement Distance** | **Maximum Movement Distance** | **Mean Movement Distance** | **Median Movement Distance** | **Maximum Movement Distance** |
| RR3 | 49.3 ± 25.9 | 50 | 100 | 50.2 ± 37.3 | 50 | 100 |
| RR10 | 35.0 ± 26.8 | 25 | 100 | 47.8 ± 37.9 | 33.3 | 100 |
| RR18 | 40.4 ± 25.0 | 50 | 100 | 19.8 ± 8.47 | 16.7 | 33.3 |
| All Sites | 39.4 ± 25.7 | 50 | 100 | 38.1 ± 32.4 | 25 | 100 |
|  | ***Erebus camprimulgus (female)*** | | | ***Erebus gemmans*** | | |
| **Site** | **Mean Movement Distance** | **Median Movement Distance** | **Maximum Movement Distance** | **Mean Movement Distance** | **Median Movement Distance** | **Maximum Movement Distance** |
| RR3 | 43.6 ± 27.9 | 33.3 | 100 | 75.0 ± 28.9 | 75 | 100 |
| RR10 | 56.9 ± 35.8 | 33.3 | 100 | NA | NA | NA |
| RR18 | 33.9 ± 15.9 | 50 | 50 | 45.0 ± 34.1 | 25 | 100 |
| All Sites | 45.1 ± 25.1 | 50 | 100 | 52.1 ± 34.6 | 50 | 100 |
|  | ***Hypoprya* spp.** | | | ***Erebus ephesperis*** | | |
| **Site** | **Mean Movement Distance** | **Median Movement Distance** | **Maximum Movement Distance** | **Mean Movement Distance** | **Median Movement Distance** | **Maximum Movement Distance** |
| RR3 | NA | NA | NA | 26.9 ± 11.8 | 25 | 50 |
| RR10 | 26.2 ± 22.2 | 18.3 | 100 | 49.0 ± 31.7 | 50 | 100 |
| RR18 | 26.3 ± 14.9 | 25 | 50 | 32.7 ± 25.8 | 25 | 100 |
| All Sites | 26.2 ± 19.2 | 20 | 100 | 34.4 ± 26.3 | 25 | 100 |

**Supplementary Figure**

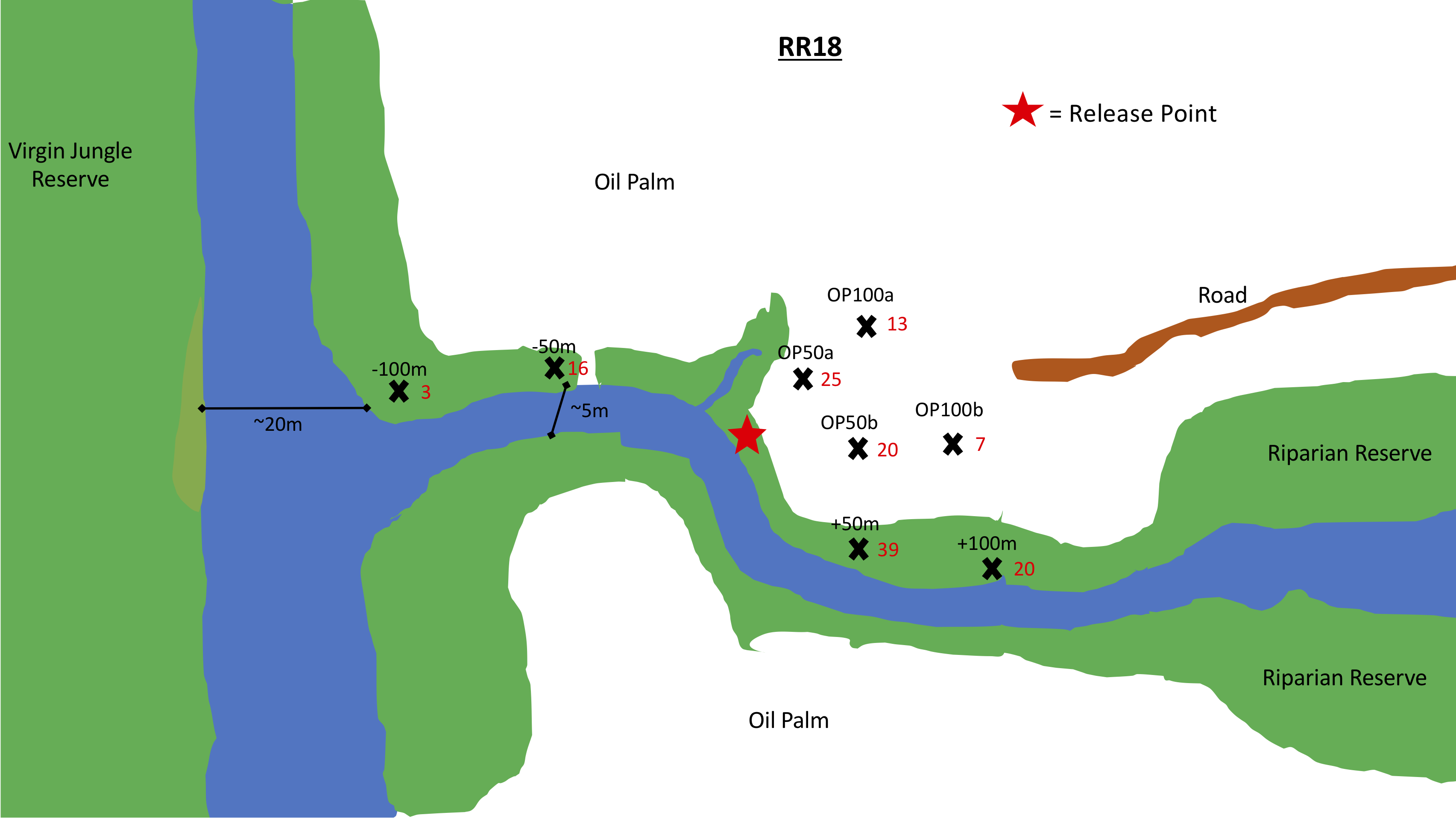

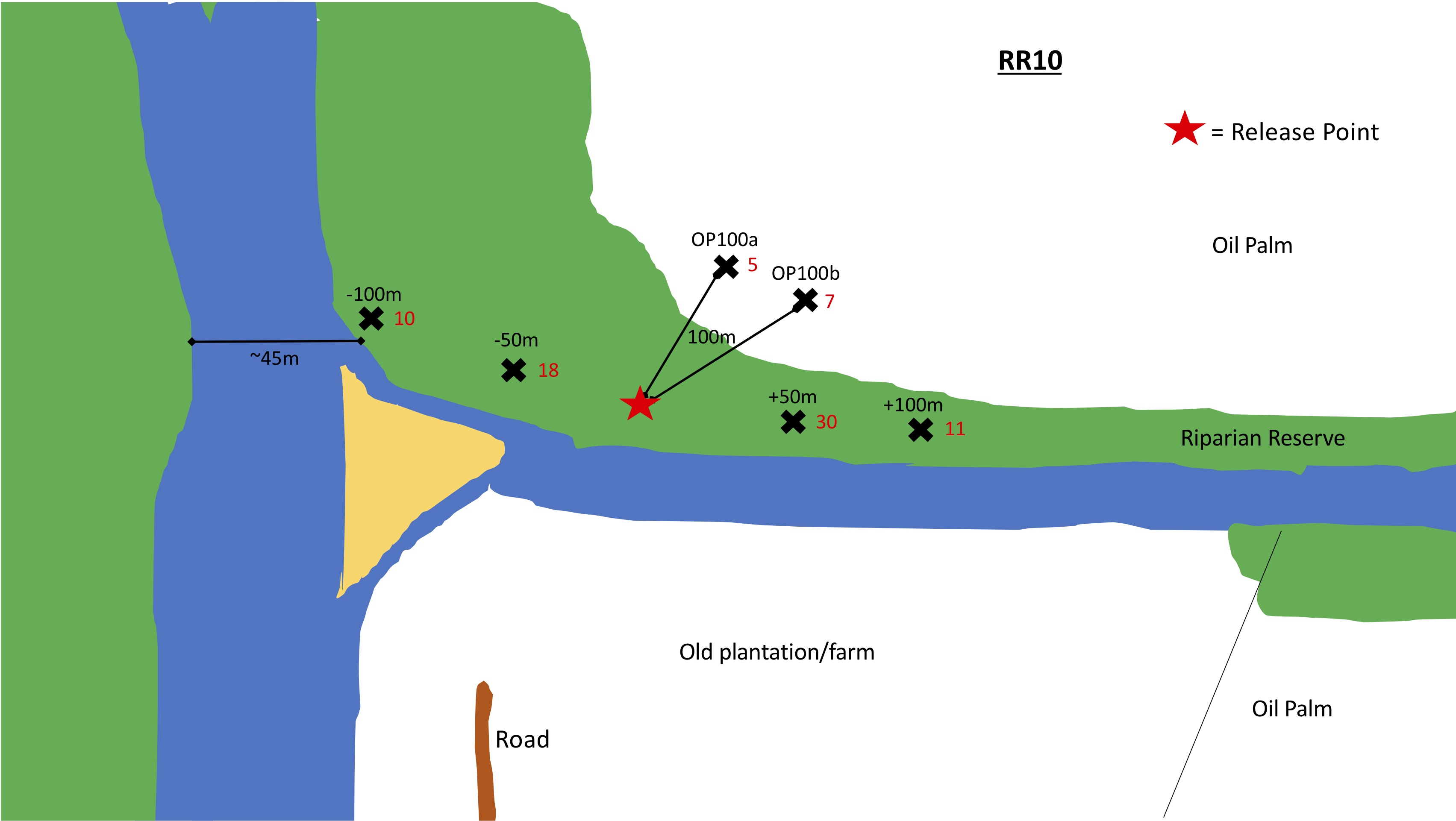

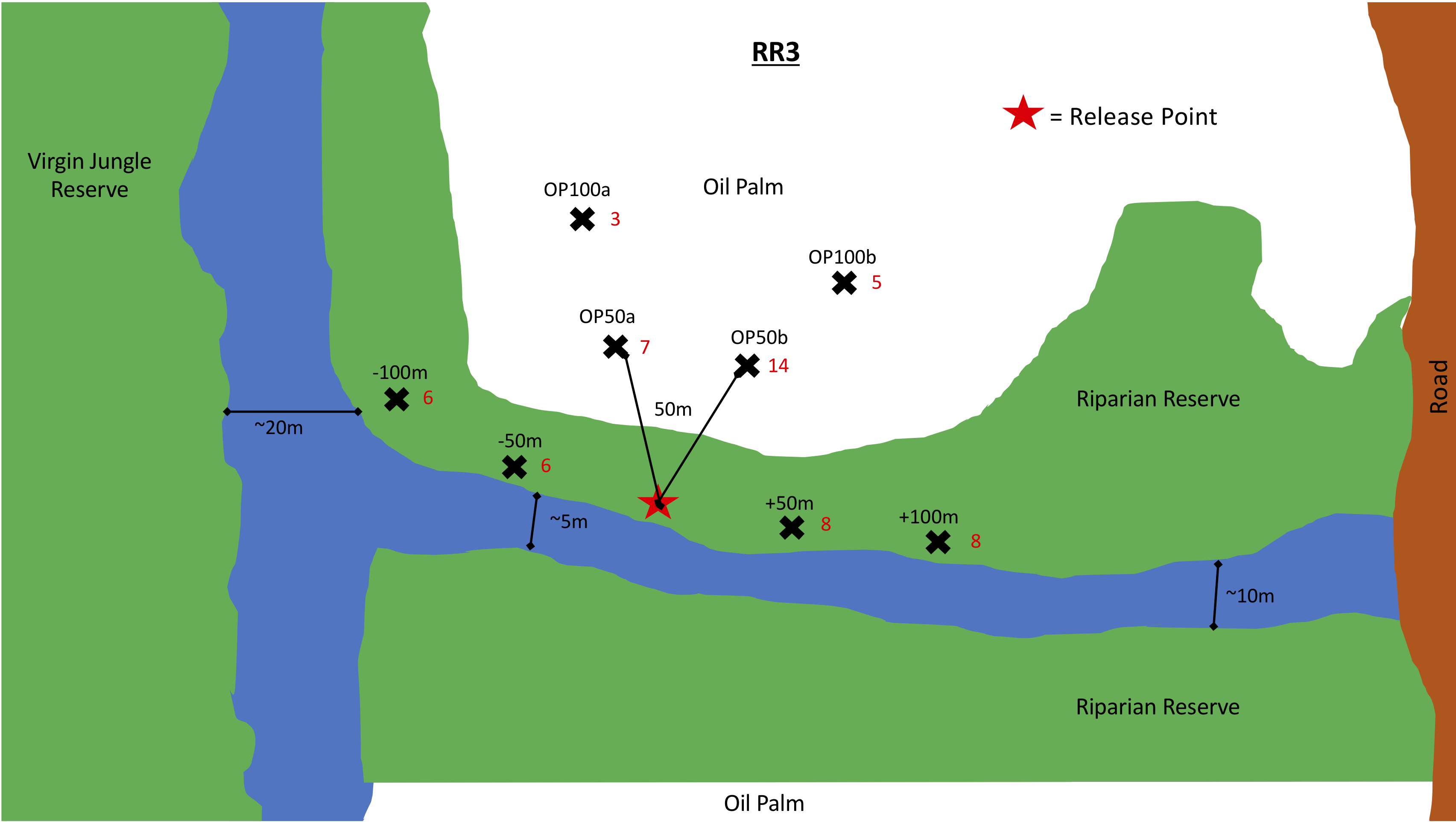

Figure S1. Site diagrams for the moth sampling. Crosses represent fruit-baited moth traps.
